# Siderophores as an iron source for *Prochlorococcus* in deep chlorophyll maximum layers of the oligotrophic ocean

**DOI:** 10.1101/2021.11.13.468467

**Authors:** Shane L. Hogle, Thomas Hackl, Randelle M. Bundy, Jiwoon Park, Brandon Satinsky, Brandon Satinsky, Teppo Hiltunen, Steven Biller, Paul M. Berube, Sallie W. Chisholm

## Abstract

*Prochlorococcus* is one of the most abundant photosynthesizing organisms in the oligotrophic oceans. Gene content variation among *Prochlorococcus* populations in separate ocean basins often mirrors the selective pressures imposed by the region’s distinct biogeochemistry. By pairing genomic datasets with trace metal concentrations from across the global ocean, we show that the genomic capacity for siderophore-mediated iron uptake is widespread in low-light adapted *Prochlorococcus* populations from iron-depleted regions of the oligotrophic Pacific and S. Atlantic oceans: *Prochlorococcus* siderophore consumers were absent in the N. Atlantic ocean (higher iron flux) but constituted up to half of all *Prochlorococcus* genomes from metagenomes in the N. Pacific (lower iron flux). *Prochlorococcus* siderophore consumers, like many other bacteria with this trait, also lack siderophore biosynthesis genes indicating that they scavenge exogenous siderophores from seawater. Statistical modeling suggests that the capacity for siderophore uptake is endemic to remote ocean regions where atmospheric iron fluxes are the smallest, particularly at deep chlorophyll maximum and primary nitrite maximum layers. We argue that abundant siderophore consumers at these two common oceanographic features could be a symptom of wider community iron stress, consistent with prior hypotheses. Our results provide a clear example of iron as a selective force driving the evolution of *Prochlorococcus*.

## Introduction

*Prochlorococcus* is the smallest known photosynthetic organism and is among the most numerically abundant life forms on the planet. It is unicellular, free-living, geographically widespread, and highly abundant in the oligotrophic subtropical/tropical ocean, often comprising half of the total chlorophyll[1]. *Prochlorococcus* and its sister lineage *Synechococcus* account for approximately 25% of global marine net primary productivity[2], making the picocyanobacteria key drivers of marine biogeochemical cycles[3]. Light gradients drive the vertical distribution of *Prochlorococcus* with low-light (LL) adapted clades occupying the deeper parts of the euphotic zone, and high-light (HL) adapted clades near the surface. This broad light-driven ecological and evolutionary division is further partitioned into mostly coherent genomic clusters (clades), each with distinct ecological and physiological properties[4]. At the finest scale of diversity, clades further partition into distinct sympatric subpopulations, which share the majority of their genes but also contain segments of unique genetic material[5]. These variable regions encode functionally adaptive traits that tune each population’s physiology to its local environment.

Iron (Fe) is a crucial micronutrient for marine phytoplankton due to its central role as an enzyme cofactor in cellular processes, including respiration and photosynthesis. As a result, the concentrations and chemical forms of Fe influence global carbon cycle dynamics[6]. Dissolved Fe (dFe) is scarce in much of the ocean and is mostly (> 99%) complexed with organic chelating ligands that solubilize and stabilize the ions in solution[7]. Concentrations of these ligands quickly increase in response to Fe fertilization events, which implies that they are actively produced by members of the microbial community[8]. It is challenging to determine the chemical structure of these ligands[9], so their abundance and stability coefficients (a measure of how “strongly” the ligand binds Fe) are typically inferred electrochemically[10]. Many electrochemical studies operationally partition the ligand pool into weak (L_2_) and strong (L_1_) binding classes. The weak L_2_ class includes humic-like substances, exopolysaccharides, and undefined colloids. The strong L_1_ class includes siderophores, small Fe-binding molecules that microbes produce during periods of Fe starvation, but it is not clear how much of the L_1_ class are genuine siderophores. Under some conditions, siderophores appear to account for most of the strong L_1_ ligand fraction in seawater[11, 12].

There are two primary mechanisms by which marine microbes extract dFe bound to the organic ligands in the ocean. First, dFe can dissociate from organic ligands in the extracellular environment (via kinetic control, photodegradation, or cell surface reductases) and is imported across the outer membrane as an unbound, inorganic ion[13]. Second, whole Fe-ligand complexes can be directly translocated across cell membranes (Fig. S1)[14]. Direct uptake pathways are prevalent in fast-growing copiotrophic marine bacteria with large genomes but absent in free-living marine bacteria with streamlined genomes such as *Prochlorococcus* or SAR11[15, 16]. In these organisms, selection favors the minimization of genome size and metabolic complexity over the versatility of maintaining multiple direct Fe uptake pathways[15]. A decade ago, it was believed that *Prochlorococcus* fulfilled its Fe requirements only via the disassociation mechanism while relying upon a single inner membrane Fe(III) ATP-binding cassette transporter[17, 18]. Prior work also showed that *Prochlorococcus* isolate genomes lacked the genes necessary for siderophore biosynthesis and uptake[18]. This image changed when putative siderophore uptake gene clusters were identified from a handful of genomes from *Prochlorococcus* surface clades HLII and HLIV collected from remote, low-Fe regions of the global ocean[16, 19]. This exciting finding suggested that some *Prochlorococcus* populations had adapted to Fe scarcity by supplementing the uptake of dissociated dFe ions with the direct uptake of siderophore-bound Fe.

Siderophore uptake genes from cells belonging to HL-adapted *Prochlorococcus* clades have been shown to be most abundant in surface waters that have low modeled Fe concentrations[20, 21]. However, extent of siderophore uptake potential in cells belonging to LL-adapted clades which are uniquely adapted to waters deep in the euphotic zone is unknown. This is important because interactions between Fe and light limitation or Fe-light co-limitation may increase Fe demand for *Prochlorococcus* clades inhabiting deeper waters and the deep chlorophyll maximum layer (DCM)[22, 23]. Also of interest is understanding what environmental features, in addition to Fe, are associated with *Prochlorococcus* siderophore uptake. Here we survey an extensive data set of picocyanobacterial uncultivated single-cell and cultivated isolate genomes to obtain a general picture of the phylogenetic and biogeographic structure of picocyanobacterial siderophore traits. Next we combine biogeochemical and metagenomic time-series datasets from the Hawai’i Ocean Time-series (HOT) station ALOHA and the Bermuda Atlantic Time-series Study (BATS) station BATS[24] to more deeply understand seasonal dynamics and the depth distributions of *Prochlorococcus* siderophore traits in a community context. Finally, we quantify the *Prochlorococcus* siderophore trait in over 700 metagenomic samples from Tara Oceans[25] and GEOTRACES[24] and connect the distribution of the trait to thousands of trace metal and other biogeochemical measurements from across the world’s oceans. Using these rich data, we identify environmental features associated with *Prochlorococcus* siderophore uptake and reveal that siderophore uptake is predominantly a feature of low-light adapted LLI clade *Prochlorococcus* residing near DCM layers from the lower limits of the euphotic zone. These findings reveal new regions of phytoplankton Fe stress in the global ocean and underscore light and Fe availability as key features shaping the evolution of *Prochlorococcus*.

## Materials and Methods

The supplementary material details all protocols and methods.

### Data sources

We used a collection of over 600 *Prochlorococcus* single-cell and isolate genomes collected from across the global ocean[26, 27], 195 surface and DCM metagenomes from Tara Oceans project[28], 480 metagenomes acquired from GEOTRACES cruises[24], and 133 metagenomes from the HOT and BATS time-series[24]. Trace metal and other chemical concentrations are from the GEOTRACES Intermediate Data Product IDP2017 version 2 (accessed January 2019)[29]. Samples were from sections GA02[30, 31], GA03, GA10[32], and GP13. Biogeochemical data from the Tara Oceans project was obtained from https://doi.pangaea.de/10.1594/PANGAEA.875579. Modeled climatological dFe from the MIT Darwin model v0.1_llc90 was obtained from the Simons Collaborative Marine Atlas project (CMAP)[33]. Modeled dFe was averaged over 12 months at a 0.5 degree grid in 5 m depth bins from the upper 250 meters. Metagenome sequence data and associated metagenomes with environmental variables were quality controlled as described earlier[34, 35].

*Prochlorococcus* cell concentrations (qPCR-calibrated) from HOT and BATS[36] were obtained from the Biological and Chemical Oceanography Data Management Office: https://www.bco-dmo.org/dataset/3381. L_1_ organic Fe-binding ligand data are from previous studies at HOT and BATS[11, 37, 38]. Siderophore concentrations from surface waters of the North Atlantic at stations 41-62 (data not available from the DCM) and the surface and DCM at HOT are from previous studies[11, 39]. The DCM was defined as the depth range of maximum chlorophyll fluorescence at HOT (100-125 m) and BATS (90-135 m). Surface was defined as all depths shallower than the DCM.

### Measurements of Fe-binding ligands and siderophores

We analyzed organic Fe-binding ligand data from BATS and HOT using competitive ligand exchange adsorptive cathodic stripping voltammetry (CLE-ACSV) as described in detail elsewhere[11, 37, 40]. We measured siderophore concentrations using liquid chromatography (LC) coupled to inductively coupled plasma mass spectrometry (ICP-MS) after pre-concentration via solid-phase extraction[11, 12, 41]. DCM samples were from a depth range of 70-200 meters, and surface samples were within a depth range of 3-45 meters.

### Comparative genomics

The steps for producing the *Prochlorococcus* genome phylogeny in Fig. 1 were detailed earlier[35]. Siderophore transporters were identified in *Prochlorococcus* genomes using the TonB dependent transporter, solute binding protein FhuD, FecCD ABC permease, ABC ATPase, TonB protein, ExbB, and ExbD proton channel families as described previously[15]. All *Prochlorococcus* genomes were searched for siderophore biosynthesis potential using antiSMASH v5.0[42], which searches for known non-ribosomal peptide synthetases and polyketide synthase siderophore biosynthesis pathways. The true proportion of genomes with the siderophore transport cluster was estimated by accounting for genome incompleteness in the single-cell genomes as described earlier[34].

**Figure 1.**
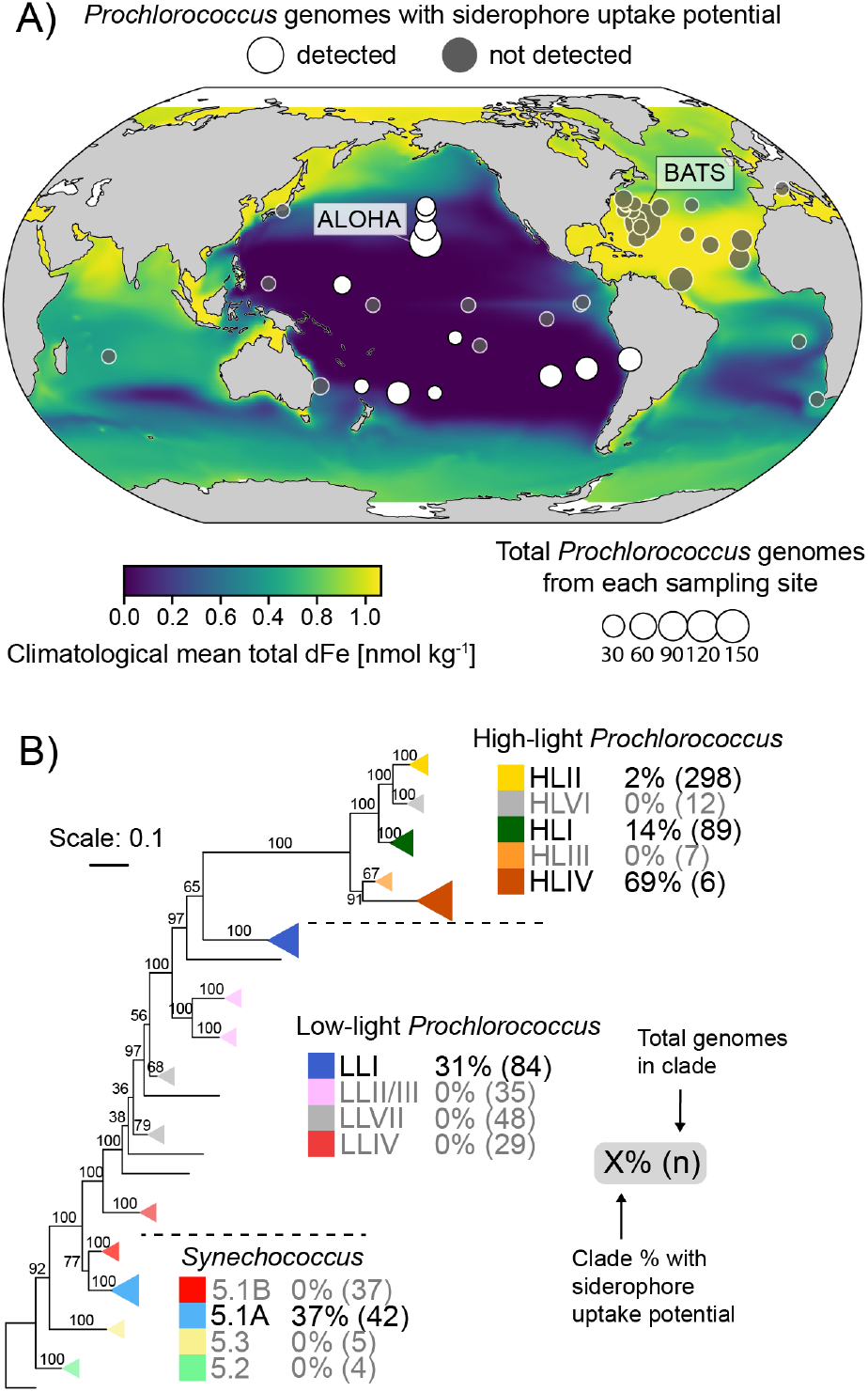
Phylogenetic and biogeographic patterns of picocyanobacterial siderophore consumers. **A)** Sampling locations of picocyanobacteria isolate and single-cell genomes used for this study. Point size is proportional to the total number of isolated genomes from each location, and point color shows whether a siderophore consumer genome was isolated at that location. Map color displays the climatological annual mean total dissolved Fe concentration nnmol kg^−1^ integrated over the upper 55 meters of the water column. Oceanographic stations ALOHA and BATS are denoted for reference. **B)** The phylogenetic tree was built with 96 *Synechococcus* genomes and 605 *Prochlorococcus* genomes and is rooted at *Synechococcus* sp. WH5701; nodes show bootstrap values (250 resamplings). Triangles depict monophyletic clades, and triangle area is proportional to the number of siderophore consumer genomes in each clade.

### Metagenomic read classification and count processing

Metagenome reads were mapped to the MARMICRODB marine microbial database using Kaiju v1.6.0[43] as described before[35]. Reads mapping best to the TonB dependent siderophore receptor or one of 730 core *Prochlorococcus* gene families were retained for further analysis. The TonB dependent receptor was selected as an indicator for the presence of the entire siderophore uptake cluster because it is the longest gene in the cluster and because its sequence composition is conserved and distinct in *Prochlorococcus* (Fig. S2). There was no clear similarity cutoff between metagenomic reads and the receptor sequence that differentiated *Prochlorococcus* clades, and we did not attempt to resolve clades by the TonB receptor (Fig. S2). Receptor and core gene counts were length-normalized to the median length of the corresponding *Prochlorococcus* gene, then divided by the median normalized abundance of the 730 core genes.

### Statistics and data analysis

Random forest regression was performed using the normalized TonB dependent receptor abundance as the response variable and a collection of 45 potential explanatory variables with the R package Ranger v0.12.1[44]. Hyperparameters were tuned and the model trained using nested ten-fold cross-validation, reserving 20% of the data for estimating final model performance. Signal extraction using principal component analysis (PCA) was performed on scaled and centered predictors[45] in cross-validation. All principal components (PCs) cumulatively explaining 99% of the variance in at least one training partition were retained (27 PCs total). Signal extraction was performed to ensure that model predictors were statistically independent and to circumvent the significant inter-variable correlations in the environmental data. PC importance rankings were determined in each training partition using the Boruta heuristic[46], resulting in all predictor PCs consistently performing better than random. This step buffers against overfitting the model while assessing variable importance relative to reference importance (i.e., noise). Boruta rankings were aggregated over the training partitions, resulting in a total of 20 PCs that, in combination, were better predictors of siderophore abundance than randomly generated data. Cumulatively, these PCs explained 97% of the variance in the original data.

After training and testing the final random forest, the original predictor variables were related to PCs using PCA loadings by taking the square of the eigenvector matrix to get the variance explained (*R*^2^) by each variable for each PC. All variables with an *R*^2^ ≥ 0.1 with at least one of the 20 informative PCs were retained (n=26). These variables were then ordered by their rank contribution to predicting the distribution of *Prochlorococcus* siderophore consumers. Rank contribution was calculated as the sum of the variance explained by each variable across all 20 informative PCs, with each PC weighted by the relative importance obtained from the feature importance algorithm. Pearson’s correlation was used to measure the strength of the linear relationship between siderophore relative abundance and each predictor variable.

Beta regression was used to model the relationship between siderophore transporter relative abundance and nitrite, DCM depth, and LLI *Prochlorococcus* abundance. LLI abundance was modeled as a constant additive effect while allowing the effect of nitrite to vary as a function of DCM depth. This model was selected because light-limited phytoplankton generate nitrite[47] and deeper DCMs are, presumably, more light-limited than shallower DCMs[48]. Continuous covariates were transformed to categorical covariates by binning into three roughly equally sized groups: 3.3e-4 < Lo NO_2_ ≤ 4.0e-2; 4.0e-2 < Md NO_2_ ≤ 7.3e-2; 7.3e-2 < Hi NO_2_ ≤ 1.4; 0.12% < Lo LLI ≤ 3.5%; 3.5% < Md LLI ≤ 9%; 9% < Hi LLI ≤ 90%. Emmeans[49] v1.6.0 R package was used to estimate the magnitude and statistical significance of marginal means for each covariate in the model. The seasonal effects in time-series metagenomes were estimated using Generalized Additive Mixed Models and Linear Mixed-Effect Models with the mgcv v1.8-26 and nlme v3.1-148 libraries in R as described earlier[34].

## Results and discussion

### Siderophore uptake potential in picocyanobacterial genomes

We searched a collection of 687 marine *Prochlorococcus* and *Synechococcus* genomes (including cultivated isolate genomes and uncultivated single-cell genomes) for siderophore biosynthesis and siderophore uptake gene clusters. These genomes are from over 40 distinct geographic locations across the world’s oceans (Fig. 1A), including the well-studied oceanographic stations ALOHA and BATS, and span the breadth of known *Prochlorococcus* and *Synechococcus* phylogenetic diversity (Fig. 1B). Samples were collected throughout the euphotic zone at depths ranging from five meters to over 200 meters depth. In agreement with previous studies, none of the *Prochlorococcus* and marine *Synechococcus* genomes contained siderophore biosynthesis gene clusters[18, 50]. However, we identified 47 genomes with siderophore transport gene clusters, hereafter siderophore consumers, (Fig. S1) and they were all either found in, or isolated from remote regions spanning the oligotrophic N. and S. Pacific Ocean gyres (Fig. 1A). *Synechococcus* siderophore consumers were from the North Pacific subtropical gyre and polar frontal regions between 28°N and 37°N[26], where surface dFe concentrations are typically low and the surface dFe inventory is depleted before the macronutrient inventory[51]. The restriction of picocyanobacterial siderophore consumers to the Pacific is analogous to the strong ocean basin segregation of genomic adaptations to nitrogen and phosphorus limitation in *Prochlorococcus[52–54]*. Because the Pacific Ocean encompasses over 70% of Fe-limited ocean regions[55] and siderophore uptake is a form of Fe scavenging[56], together these findings suggest that Fe scarcity is an important selective pressure on picocyanobacterial genome content in the oligotrophic Pacific ocean.

We next examined the distribution of the siderophore uptake trait among clades within the marine picocyanobacterial phylogeny. It is distributed unevenly among different clades of *Prochlorococcus*: 2% and 14% of the genomes from HLII and HLI clades of *Prochlorococcus*, respectively, contained these clusters, while 31% of those from LLI *Prochlorococcus* contained them (Fig. 1B, Fig. S1). The relative frequency of the siderophore trait increased significantly within *Prochlorococcus* ecotypes with lower temperature and light level optima: for example, HLI (14%, N=89) and LLI (31%, N=84)[4]. We also identified the siderophore transport gene cluster within genomes from *Synechococcus* clade 5.1A (37%, N=42) in subclades II, III, IV, UC-B, and CRD2, with subclades III and IV accounting for 70% of all *Synechococcus* siderophore consumers. Although members of *Synechococcus* clades do not stratify with depth like *Prochlorococcus* clades[57], clade 5.1A displays low-light adapted phenotypes[58] and subclades III and I (whose distribution follows clade IV) in particular are optimized for growth at low irradiance [58]. One caveat to these findings is that the total number of genomes and the clade composition between ocean basins were different so the observed gene frequencies are likely biased to some extent. Still, these findings clearly hint towards genuine trends of higher siderophore consumer frequency in the pacific ocean and in the LLI *Prochlorococcus* clade.

Previous studies of siderophore uptake genes in *Prochlorococcus* have been limited to Fe-limited surface waters which are dominated by cells belonging to HL adapted clades[19–21]. Our finding that potential siderophore uptake is most prevalent within the LLI clade (Fig. 1B), which thrives in deep waters, implies that picocyanobacterial siderophore use is most likely to be associated with low-light conditions. LLI *Prochlorococcus* is the dominant clade at DCM layers, which form near the base of the euphotic zone and are shaped by a balance between diminished light and enhanced macronutrients (N, P, Si) from deep waters. Furthermore Fe-limitation or Fe/light co-limitation is a persistent feature of this layer[22]. Phytoplankton Fe-limitation at the DCM may emerge due to the upregulation of the Fe-rich photosynthetic apparatus under low light[59], which may increase Fe demand relative to supply[60] and thus low-light *Prochlorococcus* clades likely require more Fe than high-light clades[23]. Indeed, *Prochlorococcus* has relatively high photosynthetic Fe requirements under low irradiance, and there is evidence that LLI *Prochlorococcus*, in particular, is uniquely adapted to the low-Fe and low-light conditions typical of the DCM[23]. In some cases, the high Fe requirements of LLI *Prochlorococcus* may make them more sensitive to fluctuations in Fe concentration than eukaryotic phytoplankton[61]. Competition for inorganic dFe ions at the DCM is intense[61], and the ability to utilize siderophores may allow *Prochlorococcus* to reduce relative competition for Fe through niche differentiation.

### Spatial and temporal patterns at two long-term ocean study sites

We further explored the Pacific/Atlantic divide and prevalence of the trait in LLI *Prochlorococcus* genomes by examining metagenomes from the surface, the DCM, and the bottom of the euphotic zone at Stations ALOHA and BATS. ALOHA is located in the N. Pacific subtropical gyre[62], while BATS is located in the N. Atlantic subtropical gyre. Both stations are oligotrophic, have comparable net primary productivity and carbon export[63], and are numerically dominated by *Prochlorococcus* and SAR11[64]. However, atmospheric iron fluxes are significantly higher at BATS compared with ALOHA[65]. Siderophores, strong Fe-binding ligands (L_1_), and dFe concentrations have been measured in many studies at these locations[11, 37–39]. This wealth of data allowed us to examine relationships between siderophore consumers and Fe-binding ligands, total dFe, and siderophore concentrations on seasonal time scales.

*Prochlorococcus* siderophore consumers were absent at BATS but constituted up to half of all *Prochlorococcus* genomes from DCM metagenomes at ALOHA (Fig. 2A). We could not identify one sequence similarity threshold for short metagenomic reads that distinguished *Prochlorococcus* outer membrane siderophore receptors by clade (Fig. S2). Instead, we estimated the clade-integrated proportion of siderophore consumers within the total *Prochlorococcus* population (see methods for details). The LLI clade dominates the DCM at both ALOHA and BATS[36] and has the highest frequency of single-cell genomes with siderophore transporters. It is not surprising then that the frequency of siderophore consumers in the total *Prochlorococcus* population at the DCM was strongly positively correlated with the relative abundance of the LLI clade (Fig. 2A). Siderophore consumers also followed a seasonal pattern at the ALOHA DCM, coinciding with peaks in LLI abundance in late summer and early fall (Fig. 2B) Fig. S3). This pattern was not present at BATS, where the trait was effectively absent.

**Figure 2.**
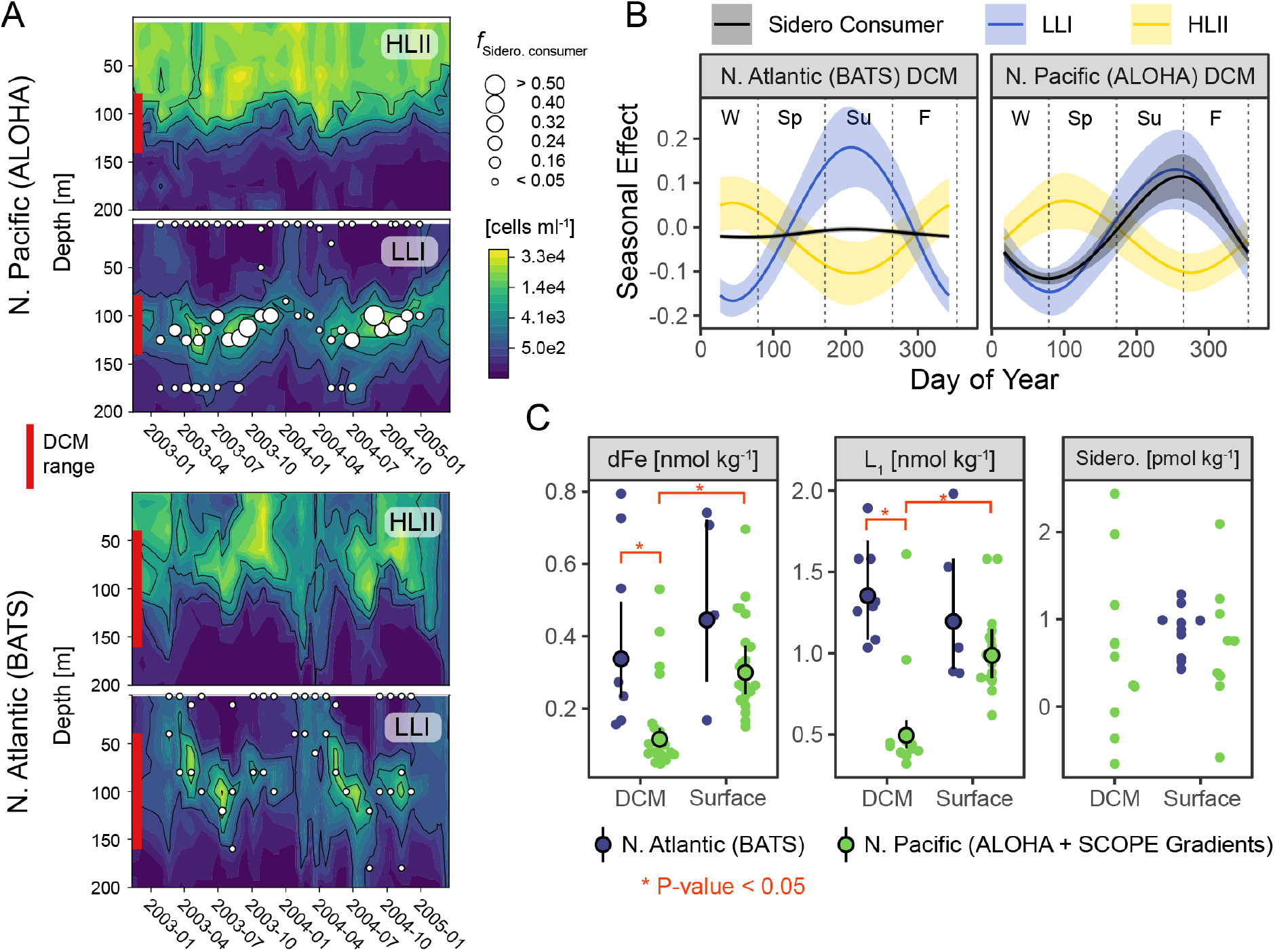
Siderophore consumers associate with the DCM in the N. Pacific, but not the N. Atlantic. **A)** *Prochlorococcus* HLII and LLI cell density over two years at HOT and BATS as determined by qPCR[36]. Contour plots of cell concentrations are cube root transformed. The deep chlorophyll maximum (DCM) depth range is highlighted in red. *f*_sidero. consumer_ is the fraction of *Prochlorococcus* siderophore consumers, and point size is proportional to relative abundance. **B)** The modeled seasonal effect (arbitrary units) on siderophore consumers, LLI, and HLII *Prochlorococcus* clades at the DCM at stations BATS and HOT. **C)** dFe concentration, L_1_ strong Fe-binding ligands, and specific siderophore classes in the N. Pacific and N. Atlantic subtropical gyres. Points and error bars are marginal population means and 95% confidence intervals from regression. Full model results are in Table S1.

We also found that biogeochemical patterns of dFe and Fe-binding ligands were inversely related to patterns of siderophore consumer abundance at stations ALOHA and BATS (Fig. 2C), Table S1). Station ALOHA had the lowest total dFe and lowest L_1_ strong ligand concentrations. The significant enrichment of siderophore consumers at the ALOHA DCM relative to the surface also coincided with a sharp decrease in total L_1_ concentration, likely due to biological demand and uptake. In short, *Prochlorococcus* siderophore consumers are only present where and when dFe and L_1_ strong ligand concentrations are lowest. The annual spring/winter Asian dust flux is the dominant new Fe source to the N. Pacific subtropical gyre[37, 51], and this annual deposition event coincides with the lowest abundance of siderophore consumers at the ALOHA DCM (Fig. 2B). Seasonal variations in Fe supply may contribute to the seasonal succession of siderophore consumers and *Prochlorococcus* clades[36] at the DCM in the subtropical N. Pacific.

### Global patterns of Prochlorococcus siderophore uptake genes

We next examined the abundance of *Prochlorococcus* siderophore consumers in ocean areas not covered by the genomic and time-series metagenomic datasets described above by quantifying the abundance of *Prochlorococcus* siderophore consumers in metagenomes from the upper 300 meters of the global tropical and subtropical ocean. In general, results from the global metagenomic data set reinforced our findings from the time-series metagenomes (Fig. 2) and the single-cell and isolate genomes (Fig. 1). Collectively, the global metagenomes revealed: 1) a high frequency of *Prochlorococcus* siderophore consumers in the subtropical and tropical Pacific relative to the N. Atlantic Ocean, and 2) high frequencies of siderophore consumers at the DCM - especially oligotrophic DCMs from the Pacific Ocean from depths over 100 meters (Fig. 3). In most Pacific DCM samples, over half of *Prochlorococcus* cells could potentially use siderophores (assuming the gene cluster is single-copy), which implies that a significant fraction of *Prochlorococcus* Fe-demand at the DCM may be fulfilled by siderophores. Unexpectedly, however, we observed a high abundance of *Prochlorococcus* siderophore transporters in the S. Atlantic subtropical gyre (Fig. 3), where the predicted climatological mean of dFe is relatively high (Fig. 1A). For example, over 40% of *Prochlorococcus* genomes were apparent siderophore consumers at the DCM in the South Atlantic ocean between 5°S and 25°S, consistent with recent studies suggesting Fe-deficiency and Fe-N co-limitation in this region[66, 67]. Indeed, there is a persistent local minimum in dFe concentrations within the euphotic zone between 5°S and 25°S on GEOTRACES transect GA02[68], and the thermocline waters (approximately 200-400 meters depth) of this region are Fe-deficient relative to macronutrients[69]. Most biogeochemical models predict community N-limitation in the S. Atlantic gyre[70, 71], but there are an increasing number of studies demonstrating Fe-limitation or Fe-macronutrient co-limitation in phytoplankton communities from this region[66, 67]. Like the equatorial Pacific, primary productivity in the S. Atlantic gyre may be primarily sustained by rapid and efficient internal Fe recycling due to low new Fe inputs[72, 73].

**Figure 3.**
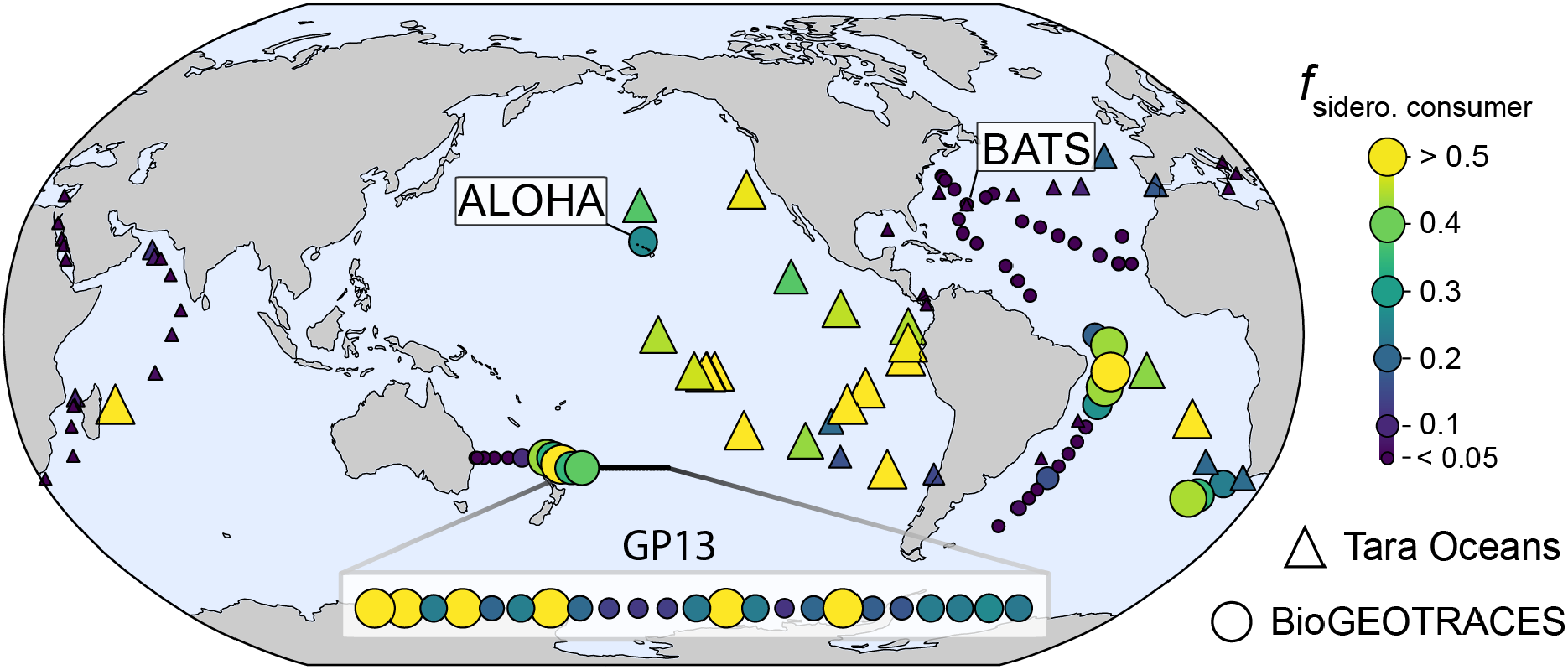
Global biogeography of *Prochlorococcus* siderophore consumers identified from metagenomes collected from deep chlorophyll maximum layers. Frequency of *Prochlorococcus* genomes with siderophore uptake gene clusters (*f*_sidero. consumer_) at the DCM in the GEOTRACES (circles) and Tara Oceans (triangles) metagenomes. Samples from the GP13 section (S. Pacific) are expanded for increased visibility. Point color and size are proportional to siderophore trait frequency.

The finding of abundant *Prochlorococcus* siderophore consumers at the DCM and in the S. Atlantic subtropical gyre led us to ask what specific chemical, biological, and hydrographic mechanisms might influence these distributions. We trained a random forest regression for *Prochlorococcus* siderophore consumer abundance using a set of 27 predictive features derived from PCA on a 45 variable dataset of hydrographic, geochemical, and biological measurements (see methods, supplementary material). We then ranked the original variables by their weighted contribution to explaining the variance in the 20 informative PCs (Fig. 4). The final model had excellent predictive performance in the test dataset (R^2^ = 0.93, RMSE = 0.00017) indicating that the abundance of *Prochlorococcus* siderophore consumers can accurately be predicted from GEOTRACES and Tara Oceans biogeochemical data. *Prochlorococcus* siderophore consumers covaried with the biogeochemical parameters that distinguish the subtropical/tropical N. Atlantic ocean, the Mediterranean Sea, and the Red Sea from the rest of the ocean. The N. Atlantic, Mediterranean Sea, and the Red Sea have the highest salinity[74] and receive a large flux of atmospherically deposited material due to their proximity to the Saharan desert and the Arabian peninsula[65]. Correspondingly, salinity and atmospherically deposited trace elements - such as Cu, AI, Zn, Mn, Fe, and Pb[75] - were strongly negatively associated with *Prochlorococcus* siderophore consumers (Fig. 4, Fig. 5, Fig. S4). Siderophore transporters were highly abundant in a handful of equatorial Pacific samples dominated by the HLIV clade (> 60% HLIV), which has been shown previously to have siderophore transport genes[19] and is associated with Fe-limited ocean regions[76]. Overall, our findings reveal that picocyanobacterial siderophore uptake is common in the remote reaches of the subtropical/tropical oligotrophic ocean, where the atmospheric input fluxes of trace elements to the upper ocean are smallest.

**Figure 4.**
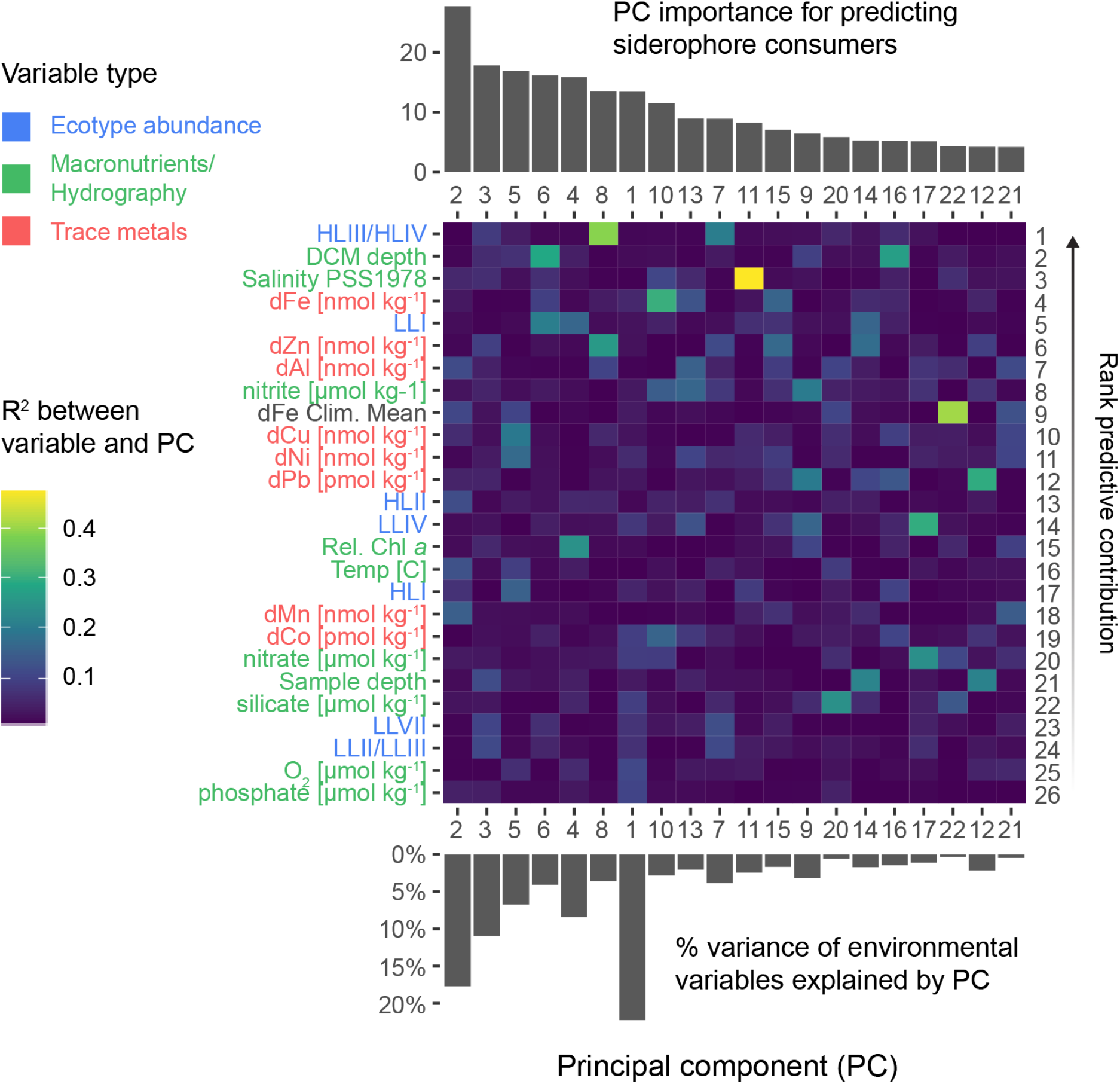
Oceanographic features associated with *Prochlorococcus* siderophore consumers. Environmental variables (heatmap left) are colored by variable type and ordered (heatmap right) by their rank contribution to predicting the distribution of *Prochlorococcus* siderophore consumers (see methods). Top barplot shows the informative principal components derived from the combined dataset and ranked by their importance (unitless) to the random forest model. The lower barplot shows the % total variance explained by each of the informative principal components. Heatmap color shows the correlation (*R*^2^) between each variable and each informative principal component.

**Figure 5.**
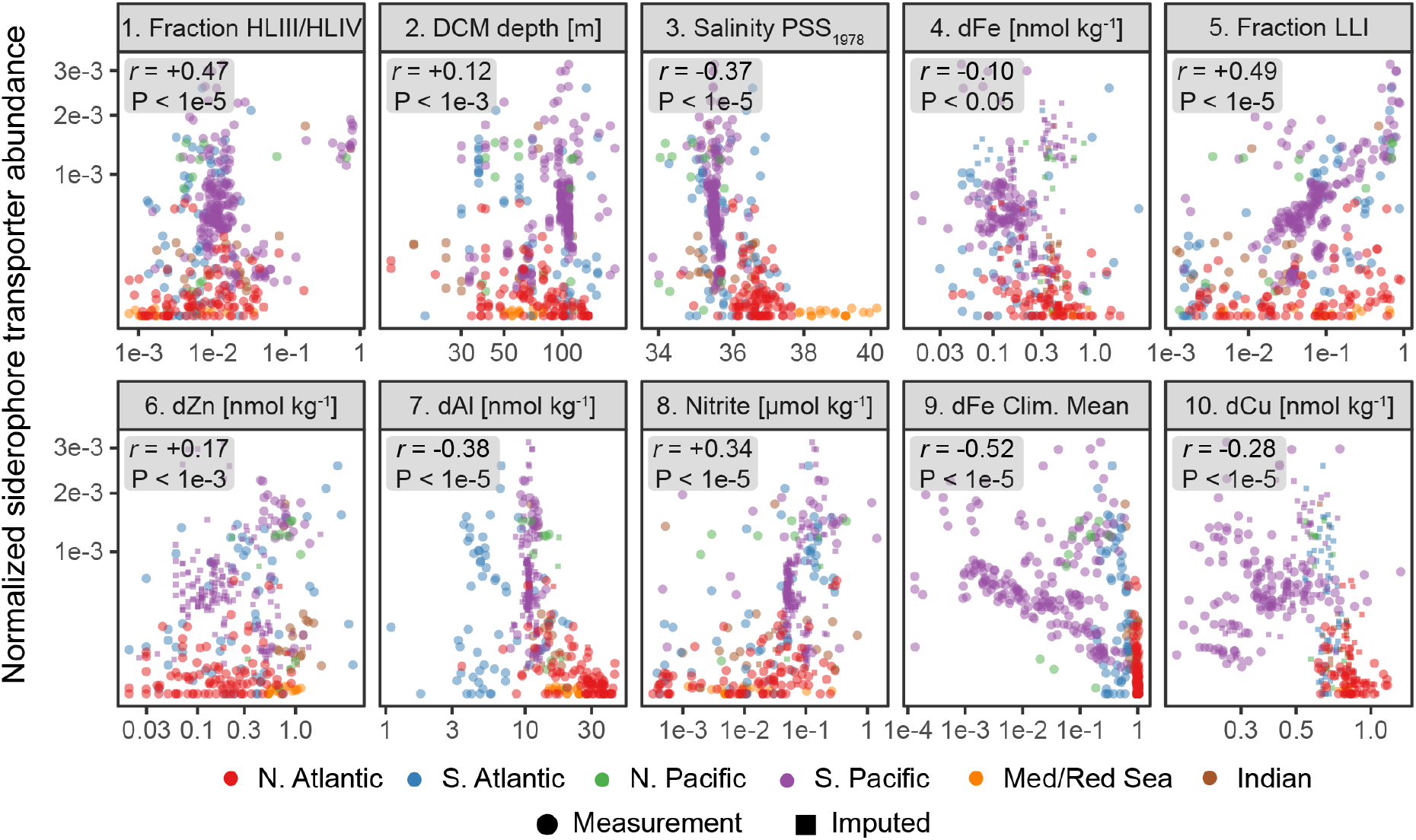
Top ten most predictive variables for *Prochlorococcus* siderophore consumer distribution. These ten environmental variables (ranked 1-10) have the greatest cumulative predictive contribution to the random forest model (Fig. 4, see methods). Each point is a metagenomic observation colored by ocean basin. Small squares represent samples with missing data that was imputed (see methods). dFe climatological means are from the MIT Darwin model. The remaining measurements are *in situ* chemical, biological, or hydrographic measurements from GEOTRACES and Tara Oceans. Boxes show Pearsons’s correlation coefficient and the p-value testing whether true *r* is equal to 0.

Of all trace metals in the global dataset, dFe measured from GEOTRACES samples explained most of the weighted variance across the principal component predictors used in the random forest. However, the magnitude of its linear correlation with siderophore consumers (*r* = −0.10) was smaller than for other metals like Al (*r* = −0.28) or Cu (*r* = −0.28) (Fig. 5). Modeled dFe concentrations from the Darwin biogeochemical model were more correlated with *Prochlorococcus* siderophore consumers (*r* = −0.54) than any trace metal measurement measurements of all trace metal measurements, including dFe. Although these variables each had a stronger linear relationship with siderophore consumers than with dFe measurements from the GEOTRACES program, dFe variance was partitioned across multiple, strongly predictive principal components (Fig. 4). This implies that dFe’s predictive power in the random forest was primarily derived from its nonlinear associations and interactions with other environmental variables (e.g., LLI abundance, DCM depth, nitrite and salinity) and not its linear correlation with siderophore consumers. This is compatible with current views of marine Fe biogeochemistry: most of the upper ocean dFe inventory is cycled rapidly and shaped by multiple interacting biotic and abiotic biogeochemical processes[73].

The strong linear correlation between modeled climatological mean dFe concentrations and *Prochlorococcus* siderophore consumers is likely because atmospheric dust deposition is the dominant Fe source to the ocean in the MIT Darwin model[77]. This mirrors the strong negative linear relationship we observe between siderophore consumers and atmospherically deposited elements (e.g., aluminum) in the GEOTRACES data and the strong partitioning of *Prochlorococcus* siderophore consumers to the oligotrophic Pacific and S. Atlantic. Thus, we argue that the global biogeography of *Prochlorococcus* siderophore consumers is driven by basin-scale geological forcing in the marine Fe cycle[75, 78]. Specifically, it appears that patterns of atmospheric dust deposition set the biogeographical boundary for where the relative fitness benefit of siderophore use exceeds that for reduced genome size and gene loss[79]. On a local scale, the fraction of a *Prochlorococcus* population that can use siderophores will also reflect a combination of physiological and ecological processes (e.g., light availability), which governs local dynamic biogeochemical processes like regeneration, scavenging, uptake, and colloid formation. In this way, the abundance of *Prochlorococcus* siderophore consumers reflects a balance of both the “fast” and “slow” biogeochemical processes shaping the marine Fe cycle.

Surprisingly, the random forest regression also showed a strong association between siderophore consumers nitrite concentrations and DCM depth. We explored this relationship further using linear regression (Fig. 6, Table S2). The results were most consistent with a scenario where *Prochlorococcus* siderophore consumers are abundant at depths with the highest nitrite concentrations, especially where the DCM layer is deeper than 100 meters and to a lesser degree where LLI cells are most abundant. We propose that this pattern is due to phytoplankton Fe and light limitation (or co-limitation) at the DCM and the primary nitrite maximum layer. The origin of the primary nitrite maximum is debated, but it likely forms due to incomplete assimilatory nitrate reduction by Fe- or light-limited phytoplankton[48] in parallel with uncoupled chemoautotrophic nitrification[80]. Due to fast phytoplankton uptake kinetics, nitrite should only accumulate in waters with low photon flux, where phytoplankton inorganic nitrogen uptake is limited by Fe and light[80]. Nearly all LLI cells can assimilate nitrite and some can also assimilate nitrate[54], but we found no evidence that nitrate assimilation genes were more or less abundant in LLI siderophore consumers than expected by chance. Overall, the significant positive association of siderophore consumers and LLI *Prochlorococcus* with high nitrite samples - especially with the deepest DCMs - provides indirect support for the hypothesis that the primary nitrite maximum layer in the euphotic subtropical/tropical ocean is associated with Fe and light limitation of photoautotrophic cells.

**Figure 6.**
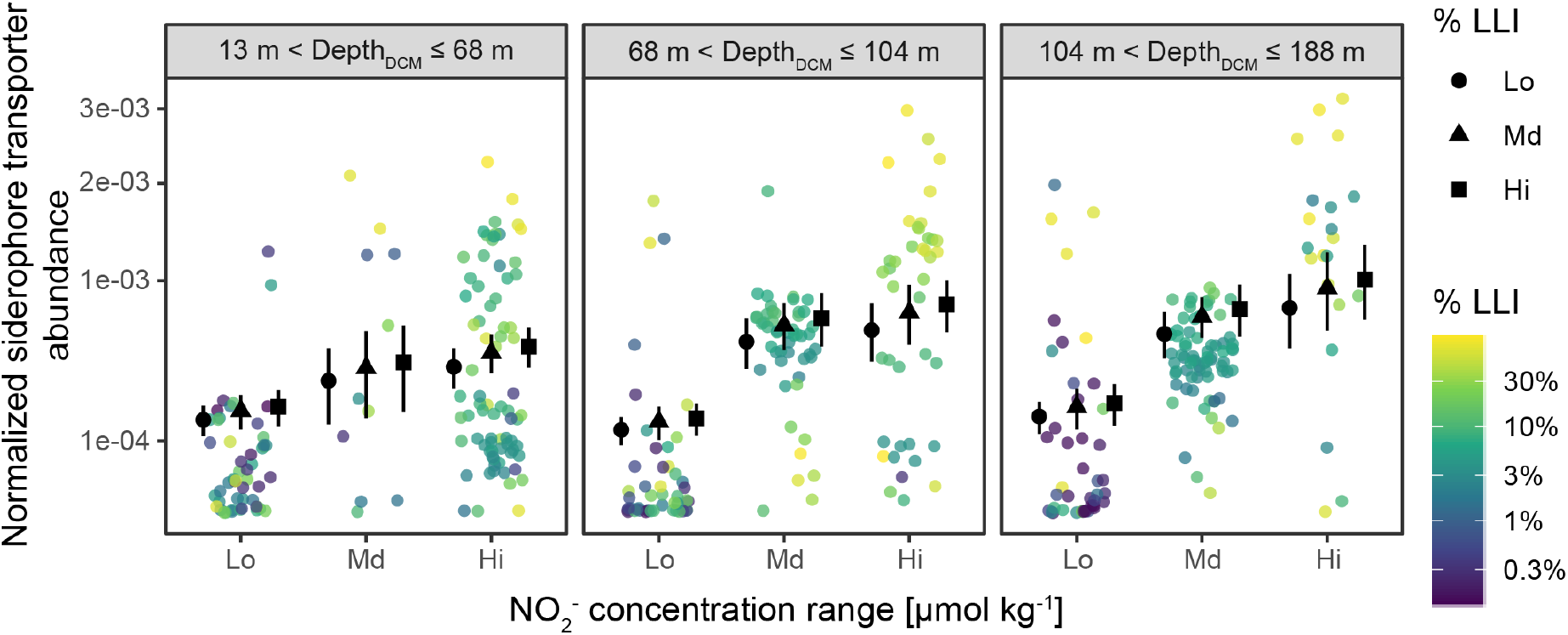
*Prochlorococcus* siderophore consumers associate with the highest nitrite concentrations and the deepest deep chlorophyll maximum layers. Each point is a direct metagenomic observation colored by the percentage of LLI clade in each sample. Subplots show different DCM depth ranges. Points and error bars show estimated marginal population means and 95% confidence intervals from beta regression using LLI % clade abundance, nitrite concentration, and DCM depth as model covariates. These continuous covariates were transformed into categorical covariates for regression by binning into three equal-sized groups (see methods).

## Synthesis

High-frequency dynamics in the organic Fe-binding ligand pool of the upper ocean balance the interplay between regeneration and removal fluxes which determines the global inventory of oceanic Fe[73, 81]. Siderophores are increasingly understood to be an important molecular mechanism regenerating Fe from biomass and incorporating atmospherically deposited lithogenic Fe into microbial ecosystems[56]. Many copiotrophic marine bacteria produce and use siderophores, but nearly all genome-streamlined oligotrophic marine bacteria do not - likely to minimize overall metabolic complexity[15, 16]. This raises the question of how minimalist cells like *Prochlorococcus* fulfill their Fe requirements, especially in the most Fe-limited regions.

Using rich genomic and metagenome data sets, we found that large populations of *Prochlorococcus* and *Synechococcus* from remote regions of the global ocean have evolved the ability to scavenge exogenous siderophore-bound Fe, which likely helps them fulfill cellular iron demand in limiting or stressful conditions. The siderophore uptake trait is prominent in ocean regions where the atmospheric Fe flux to the surface ocean is lowest (Fig. 7). In these regions, Fe recycling mediated by organic ligands fuels the majority of primary productivity[72]. Here siderophores are likely a critical molecular shuttle between particulate, colloidal, and dissolved phases, ultimately retaining Fe in the upper ocean (Fig. 7). The siderophore uptake trait is most abundant in picocyanobacteria inhabiting remote Fe-depleted regions and in low-light adapted *Prochlorococcus*, particularly those at deep DCMs near the primary nitrite maximum layer. The vertical distribution of this trait in the water column implies that light availability is also an important control on *Prochlorococcus* Fe demand. Indeed, the association of siderophore consumers with the highest nitrite concentrations suggests that nitrite accumulation in the euphotic zone may be a consequence of both iron and light limitation.

**Figure 7.**
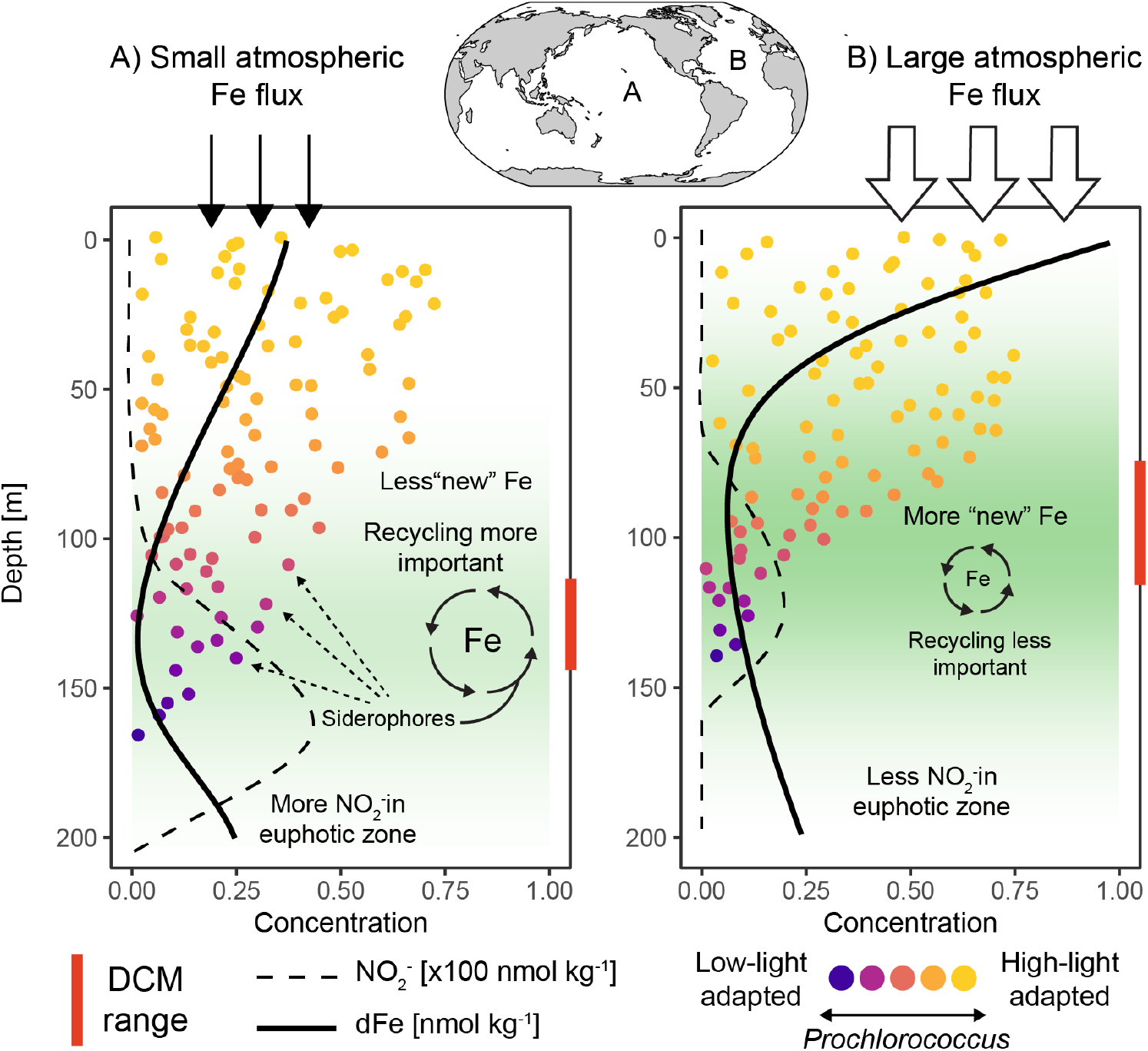
Biogeochemical drivers of *Prochlorococcus* siderophore uptake. Differences in picocyanobacteria siderophore use between the Pacific and Atlantic ocean. The total abundance of *Prochlorococcus* is proportional to the number of points at each depth, while color represents high-light and low-light adaptation. DCM = deep chlorophyll maximum layer.

Our results imply that the presence of siderophore-consuming *Prochlorococcus* could be a valuable biomarker for diagnosing ecosystem Fe deficiency. Indeed, our study highlights two areas of potential iron-stress that warrant further investigation - the S. Atlantic subtropical gyre and the DCM of the oligotrophic Pacific and S. Atlantic oceans. These areas have historically been considered nitrogen- and light-limited ecosystems, but our results suggest an essential role for Fe. Most of the potentially Fe-deficient regions revealed by our analysis are remote, and sampling Fe at these locations requires specialized knowledge of shipboard trace metal incubations, sampling, and analysis. Because DNA sampling does not require specialized trace metal clean conditions, omics-based approaches based on robust markers of nutrient deficiency could be one complementary tool for increasing the breadth of trace metal surveys in the ocean. Here we have leveraged cross-scale biology of the well-described and ubiquitous marine picocyanobacterium, *Prochlorococcus*, to reveal vast regions of community iron deficiency in the global ocean using a siderophore transport system as a biomarker. These findings highlight the intricate connection between the ecology and evolution of *Prochlorococcus* and the Fe cycle of the surface ocean.

## Data and materials availability

GEOTRACES and HOT/BATS Metagenome sequencing reads are available from the NCBI Sequence Read Archive under studies SRP110813 and SRP109831 and BioProjects PRJNA385854 and PRJNA385855. Associated sample collection data can be found at https://dx.doi.org/10.1038/sdata.2018.176. Tara Oceans Primary Metagenome sequencing reads are available from the European Nucleotide Archive under project PRJEB402. Associated Tara Oceans chemical and environmental data are available from the Pangaea repository at https://doi.pangaea.de/10.1594/PANGAEA.875576. Chemical data from the GEOTRACES Intermediate Data Project IDP17 v2 are available at https://www.bodc.ac.uk/geotraces/data/idp2017/. Picocyanobacterial single cell genomes and other sequence data are available from GenBank 16S/ITS records MG666579-MG668595, MH074888-MH077527, MH319718-MH319767, and MH327275-MH327492; NCBI Sequence Read Archive study SRP141175; and Genbank assemblies QBVH00000000-QCVZ00000000. Associated sample collection data can be found at https://dx.doi.org/10.1038/sdata.2018.154. Output from the MIT Darwin model and qPCR measurements of *Prochlorococcus* abundance at HOT/BATS is available from the Simons Collaborative Marine Atlas Project https://simonscmap.com/. The MARMICRODB reference database and instructions for use are available from https://dx.doi.org/10.5281/zenodo.3520509. All computer code for reproducing the results from this specific study is available from https://github.com/slhogle/cyano-sidero-ocean.

## Acknowledgments

We thank present and past members of the GEOTRACES consortium for collecting, curating, and sharing trace metal and other biogeochemical data. In particular, we thank the GEOTRACES chief scientists for their support: Andrew Bowie (University of Tasmania), Philip Boyd (University of Tasmania), Edward Boyle (Massachusetts Institute of Technology), Gregory Cutter (Old Dominion University), Loes Gerringa (NIOZ Royal Netherlands Institute for Sea Research), Gideon Henderson (University of Oxford), William Jenkins (Woods Hole Oceanographic Institution), and Micha Rijkenberg (NIOZ Royal Netherlands Institute for Sea Research), as well as Gerhard Herndl (University of Vienna), James Moffett (University of Southern California) and Hein de Baar (NIOZ Royal Netherlands Institute for Sea Research) for their support with GEOTRACES. We thank the HOT and BATS field teams and organizational leaders for their assistance. We also thank the developers, maintainers, and administrators of the Simons Collaborative Marine Atlas Project and the MIT Darwin ecosystem model. This work was supported in part by grants from the National Science Foundation (OCE-1153588, and DBI-0424599 to SWC) and the Simons Foundation (Life Sciences Project Award IDs 337262, 647135 to SWC; SCOPE Award ID 329108 to SWC. This paper is a contribution from the Simons Collaboration on Ocean Processes and Ecology (SCOPE).

## Author contributions

SLH - Conceptualization, Investigation, Data curation, Software, Formal analysis, Visualization, Writing original draft, Writing review and editing; THackl - Investigation, Data curation, Software, Formal analysis, Writing review and editing; RMB - Investigation, Data curation, Methodology, Writing review and editing; JP - Data curation, Methodology, BMS - Data curation, Writing review and editing, THiltunen - Resources, Investigation, Writing review and editing; SJB - Data curation, Investigation, Writing review and editing; PB - Data curation, Investigation, Writing review and editing; SWC - Conceptualization, Supervision, Project administration, Resources, Funding acquisition, Writing review and editing.

