## Supplementary material for "Siderophores as an iron source for *Prochlorococcus* in deep chlorophyll maximum layers of the oligotrophic ocean"

This PDF file includes:
Supplementary Text
Supplementary Figures S1 to S4
Supplementary Tables S1 and S2

### 1 Supplementary Methods

#### 1.1 Data sources

The collection of genomes searched in this study (MARMICRODB) was previously introduced as a reference database for quantifying *Prochlorococcus* and SAR11 abundance in marine metagenomes[1, 2]. These genomes were selected to provide a comprehensive representation of the marine environment while also affording a nonredundant and simplified characterization of terrestrial and host-associated systems. MARMICRODB includes 18758 bacterial and archaeal single cell, metagenome-assembled, and isolate genomes and eukaryotic transcriptomes from a variety of sources[3–12] as well as over 7000 viral genomes from NCBI RefSeq[13].

The metagenomic collection consisted of 195 surface and deep chlorophyll maximum (DCM) metagenomes from Tara Oceans project[14] and 480 metagenomes acquired from GEOTRACES cruises[15]. *Prochlorococcus* clade abundances from GEOTRACES metagenomes were obtained from a previous study[1]. *Prochlorococcus* clade cell concentrations from the Hawai'i Ocean Time-series Station ALOHA and the Bermuda-Atlantic Time-series Study BATS site were accessed from the Biological and Chemical Oceanography Data Management Office (`https://www.bco-dmo.org/dataset/3381`). Clades were measured by quantitative PCR as described in a prior study[16].

Trace metal and other chemical concentrations were obtained from the GEOTRACES Intermediate Data Product IDP2017 version 2 (accessed January 2019)[17]. Samples were included from sections GA02[18, 19], GA03, GA10[20], and GP13, which all had paired metagenomic data. Biogeochemical data from the Tara Oceans project[21] was downloaded from `https://doi.pangaea.de/10.1594/PANGAEA.875579`. Climatological mean dissolved iron from the MIT Darwin model (v0.1_llc90, `http://darwinproject.mit.edu/`) was obtained using pycmap v0.1.2 [22]. This model version is modified from earlier versions[23, 24] and includes biogeochemical cycling of nutrient elements and a planktonic marine ecosystem model incorporating functional and size diversity with multiple links between trophic levels. It is driven by the physical ocean circulation model from ECCO v4[1]. The iron cycle parameterization in the MIT Darwin model includes a dominant iron source through aeolian dust deposition and a dominant sink due to particle scavenging and dissolution/complexation with organic ligands. The model includes two chemical species - $Fe'$ (inorganic free ferric iron) and FeL (organically complexed Fe). The total dissolved iron concentration consists of the sum of these two species.

#### 1.2 Preprocessing biogeochemical and hydrographic measurements

Raw fluorescence or derived chlorophyll *a* concentrations from hydrographic CTD casts on the GEOTRACES cruises were obtained from the IDP17[17]. At each sampling station, the fluorescence signal from the CTD profile was smoothed using a 10-meter rolling mean, then scaled at each depth by the profile standard deviation, and finally divided by the maximum value to obtain a normalized and scaled fluorescence profile from each sampling location. Normalization was necessary because fluorometer data from across the datasets was either in uncalibrated factory units, derived chlorophyll concentrations, or raw signal and individual instrument calibration coefficients were unavailable. This normalization procedure discards the fluorescence signal magnitude, but it allows for a between-sample comparison of fluorescence profile shape. The DCM depth was defined as depths with fluorescence values within 90% of the peak subsurface fluorescence signal and contiguous to the maximum fluorescence depth. The full DCM depth range was defined as 50% of maximal subsurface fluorescence.

When metagenome bottle samples did not align precisely with the depth/coordinates of the hydrographic casts or trace metal casts, the nearest CTD cast (within a 100 km radius) was matched to discrete bottle data from within a 15-meter depth range. Coordinates were matched using the shortest distance between two geospatial points according to the Haversine formula which assumes a spherical earth and ignores ellipsoidal effects. Despite this relaxed, fuzzy-matching approach, some combinations of predictor covariates were missing for some observations in the dataset. Generally, features like salinity, temperature, and macronutrient concentrations were present for most metagenome observations. However, in fewer than 5% of all samples, we needed to impute at least one hydrographic parameter or macronutrient covariate. We imputed missing salinity, temperature, macronutrient, and DCM depth observations by the missForest algorithm [25] using chaining random forest prediction and imputation from all other available variables except siderophore transporter abundance. This process was implemented with the ranger package [26] in missRanger v2.1.3 (`https://github.com/mayer79/missRanger`). The GEOTRACES trace metal data was not as complete as the macronutrient and hydrographic data. For example, measured dissolved iron or copper concentrations from GEOTRACES IDP17 were only available for roughly half of the total metagenome observations. Therefore, we also used random forest imputation for trace metal measurements, but we only imputed trace metal categories with no less than 50% missing measurements. We only imputed missing samples using data from the same ocean basin (N. Atlantic, S. Atlantic, N. Pacific, S Pacific, Mediterranean and Red Seas, and Indian) to maximize the chance of preserving basin-scale biogeochemical differences in metal concentrations. The imputation workflow is documented at `https://github.com/slhogle/cyano-sidero-ocean`.

#### 1.3 Homology searches in genomes

*Prochlorococcus* (N=663) and *Synechococcus* (N=96) genomes from MARMICRODB were searched for the trace metal uptake families following earlier studies[27] using HMMERv3.1b2 (`http://hmmer.org/`) (hmmsearch, hmmscan). Searches were focused on a selection of families related to siderophore transport, including the TonB dependent transporter (Pfam ID: PF00593, COG ID: COG4771), solute binding protein FhuD (Pfam ID: PF01497, COG ID: COG0614), FecCD ABC permease (Pfam ID: PF01032, COG ID: COG0609), ABC ATPase (Pfam ID: PF00005, COG ID: COG1120), tonB protein (Pfam ID: PF03544, COG ID: COG0810), ExbB proton channel family (Pfam ID: PF01618, COG ID: COG0811), and the biopolymer transport protein family ExbD (Pfam ID: PF02472). Based on the presence of these protein families, 33 *Prochlorococcus* genomes and 13 *Synechococcus* genomes with putative siderophore uptake clusters were identified. Pairwise sequence similarity between TonB dependent receptors from MARMICRODB were determined using blastp with default parameters[28].

#### 1.4 Estimation of siderophore transport cluster prevalence in genomes

Many of the analyzed *Prochlorococcus* and *Synechococcus* genomes were incomplete because they were sequenced from material isolated from single cells. Therefore, genome completeness was taken into account when estimating the proportion of genomes with the siderophore transport cluster. It was assumed that there was effectively no bias in the content of genome recovery in the single-cell genome sequencing and assembly processes, which is reasonable for the latest single-cell technologies[29]. The number of missing base pairs in each incomplete assembly was estimated with checkM using a custom set of 730 single-copy, core genes from closed, isolate *Prochlorococcus* genomes [30, 31]. The CheckM taxonomic-specific workflow was used for the genus *Synechococcus*. The corrected prevalence of the tonB dependent transporter gene was estimated as:

$$f = \frac{\sum_{i=1}^{n} p_i}{\sum_{i=1}^{n} g_i} \times \sum_{i=1}^{n} \left(g_i/c_i\right)/\bar{p}/n$$

where for each taxonomic group/clade, $p$ is the length of the tonB dependent transporter in base pairs, $g$ is the length of the genome assembly in base pairs, $c$ is the completeness estimate, $\bar{p}$ is the average length of all tonB dependent transporters from each taxonomic group/clade, and $n$ is the total number of assemblies from each taxonomic group/clade. The tonB dependent transporter gene always co-occurred with other transport components (ABC ATPase, ABC permease, ABC solute binding protein, TonB, ExbB, ExbD) as an operon-like structure within the picocyanobacterial genomes. Hence, it was assumed that the corrected prevalence of this individual gene represents that of the entire siderophore transport cluster.

#### 1.5 Quality control of metagenome sequencing reads

Raw metagenomic reads were preprocessed using the bbtools software suite (`https://jgi.doe.gov/data-and-tools/bbtools/`) as described in detail earlier[1]. Briefly, reads were trimmed of sequencing adapters using kmer matches to Illumina adapter sequences from the 3' end of the read. A maximum Hamming distance of one was required for matches to reference kmers. Reads with three or more 'Ns' or with an average quality score of less than Q20 were discarded. Overlapping reads were trimmed based on insert size (tbo=t) if adapter kmers were not identified and then both reads trimmed to the minimum length of the read pair. Reads shorter than 60 bp after all adapter trimming steps were discarded. Reads were also discarded if they contained common Illumina sequencing artifacts (based on kmer matches, k=31). Reads were overlapped using the bbmerge tool[32] using default parameters with a minimum insert size of 35 bp and a minimum overlapping sequence of 12 bp. All unmerged and orphaned reads that passed quality control were also retained for downstream annotation and analysis.

#### 1.6 Metagenome read classification

Metagenome reads were mapped to MARMICRODB using Kaiju v1.6.0[33] as described before[1]. Briefly, Kaiju was run in greedy mode, tolerating a maximum of five amino acid mismatches per match, a minimum match length of 11, a minimum BLOSUM62 score of 65, and excluded alignments with a Kaiju "expect value" (analogous to BLAST e value) less than 0.05. Query sequences with low-complexity regions were filutered using the SEG filtering algorithm from the BLAST+ package[28]. The majority of reads (54

#### 1.7 Processing and analysis of metagenomic count data

Sequence identifiers and taxonomic assignments for each read were parsed from Kaiju tabular output. For each sample, reads with a best match to any *Prochlorococcus* TonB dependent receptor were summed then length-normalized (reads per kilobase; RPK) to the median length of *Prochlorococcus* tonB receptors. A collection of 730 single-copy core genes from 623 *Prochlorococcus* genomes (`https://doi.org/10.5281/zenodo.3719132`) was used to estimate the abundance of *Prochlorococcus* genomes in each sample. For each sample, reads mapping to each core family were length-normalized to the median length of the family. Samples where the median *Prochlorococcus* marker gene coverage was less than 100 RPK were excluded. Outlier core gene families were excluded if they were not present in all samples and were outside the 10th and 90th RPK percentiles of all markers across all samples. The outlier removal process resulted in 277 robust *Prochlorococcus* core marker gene families that were stable across all samples (i.e. had no significant differences between ocean basins or depth ranges). The abundance of *Prochlorococcus* genomes in each sample was taken as the median of the 277 length-normalized core gene abundances. The frequency of siderophore consumers was calculated as the ratio of length-normalized TonB outer membrane receptor abundance to the length-normalized abundance of *Prochlorococcus* genomes. Assuming that siderophore transporters are single-copy per genome, this value should approximate the proportion of *Prochlorococcus* genomes in a metagenomic sample with the potential to consume siderophores.

#### 1.8 Random Forest training and testing

A random forest regression was trained to predict the frequency of *Prochlorococcus* siderophore consumers using 45 different environmental features as input. These environmental features included trace metal data from GEOTRACES, macronutrient data from GEOTRACES and Tara Oceans, modeled climatological dFe means from the MIT Darwin model (`http://darwinproject.mit.edu/`), frequencies of different *Prochlorococcus* ecotypes, and water column chlorophyll a profiles. Samples were restricted to the upper 300 meters of the water column and where the median RPK of marker genes was higher than 100 (n = 393 metagenomic samples). Random forest hyperparameters were tuned using a grid of potential parameter combinations derived from latin hypercube sampling. The latin hypercube algorithm attempts to maximize the determinant of the spatial correlation matrix between coordinates in parameter space. Model training was performed using nested ten-fold cross-validation, reserving 20% of the data for estimating final model performance. Specifically, the number of variables to split at each tree node (mtry), the minimum number of observations to partition at each node (min.node.size), and the forest size (ntrees) were tuned. Hyperparameters were selected by maximizing the coefficient of determination from correlation (model $R^2$) and minimizing the root mean square error (RMSE). Performance metrics were averaged over the ten cross-validation splits of the training data.

Random forest regression internally ranks predictor variables by their relative importance for completing a given prediction task, but it can not by itself assess whether the influence of a variable is greater than expected by chance under a given probability distribution. The latter task is necessary, for example, if one is interested in understanding the mechanisms governing a system rather than the simple construction of a “black-box” predictive tool. The Boruta heuristic was used to assess variable importance with 100 random permutations for feature selection and importance ranking[34]. Briefly, Boruta is designed to identify all dataset features more relevant to a random forest regression task than by chance alone[34]. In practice, Boruta will identify variables that may be redundant/correlated or that weakly interact with other features to produce a correlation with the outcome variable greater than that of random noise. During one algorithm iteration, Boruta doubles each feature in a dataset, randomly shuffles the doubled variables’ observed values, and constructs a random forest model with the original and randomized features. Then by using the precomputed importance rankings from the model construction step, Bortua compares the importance of the ‘true’ features to the maximum importance of the randomly shuffled ‘shadow’ features using a two-sided test of equality (p-value cutoff of 0.01). This process is then repeated 100 times or until all features have a consistent designation of importance. By iterating this process, the algorithm identifies dataset features that consistently underperform or overperform relative to chance.

#### 1.9 Dataset and code availability

The data and code for generating all figures and tables and supporting the text’s conclusions are available from `https://github.com/slhogle/cyano-sidero-ocean`.

The MARMICRODB dataset, including a comprehensive description, raw protein sequence files, Kaiju formatted databases, scripts, and instructions for using the resource, is available from `https://doi.org/10.5281/zenodo.3520509`.

The list of *Prochlorococcus* core PFAM and TIGRFAM families, a compiled HMMERv3 hidden Markov model database, and a CheckM formatted marker list file are available from `https://doi.org/10.5281/zenodo.3719132`.

### Supplementary Figures: 173

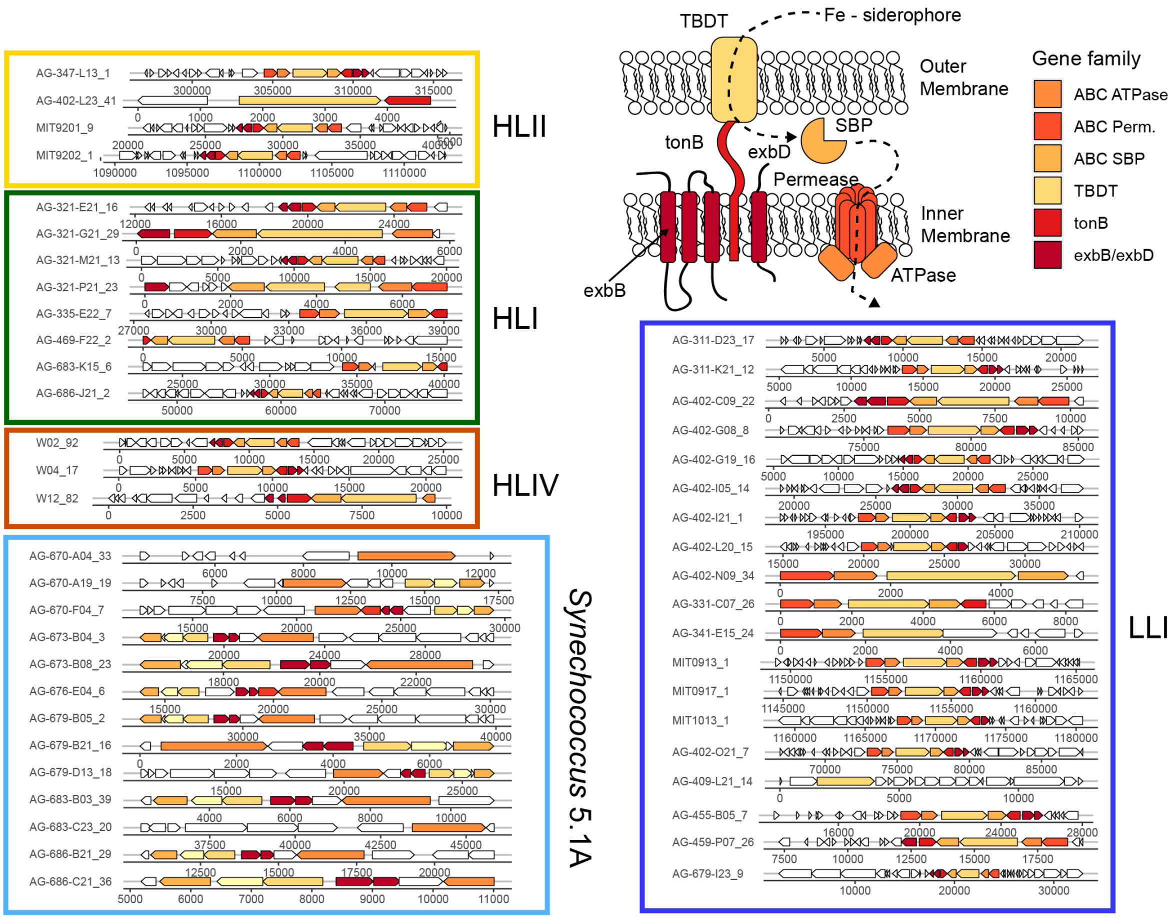

**Supplementary Figure S1.** Gene diagrams of siderophore transport clusters in marine picocyanobacteria Gram-negative bacteria import Fe-bound siderophores in a multistep process involving three protein modules: 1) an outer membrane, tonB-dependent transporter (TBDT) that recognizes and binds loaded siderophores, 2) an energy transducing complex (TonB/ExbB/ExbD) that translocates siderophores through the TBDT via proton motive force, and 3) an ATP-binding cassette (ABC) complex consisting of an inner membrane-bound permease/ATPase complex and a periplasmic solute binding protein[35]. Bacteria synthesize siderophores from secondary metabolite gene clusters encoding nonribosomal peptide synthetases and polyketide synthases[36]. The genes encoding both siderophore biosynthesis and uptake are typically clustered on the chromosome in an operon-like structure and are tightly regulated. Here gene diagrams show the absolute position of siderophore transport components and a cartoon diagram of the TonB-dependent transport components for siderophore uptake. Diagrams are partitioned by ecotype/clade (colored boxes). The last number following each genome is the contig number of the gene cluster. ABC ATPase, ATP-binding cassette transporter ATPase component (COG1120); ABC Perm., ATP-binding cassette transporter membrane permease component (COG0609); ABC SBP, ATP-binding cassette transporter periplasmic solute binding component (COG0614); TBDT, outer membrane TonB-dependent receptor protein (COG4771); tonB, energy transducing protein TonB (COG0810); exbB/exbD, biopolymer transport proteins (COG0811).

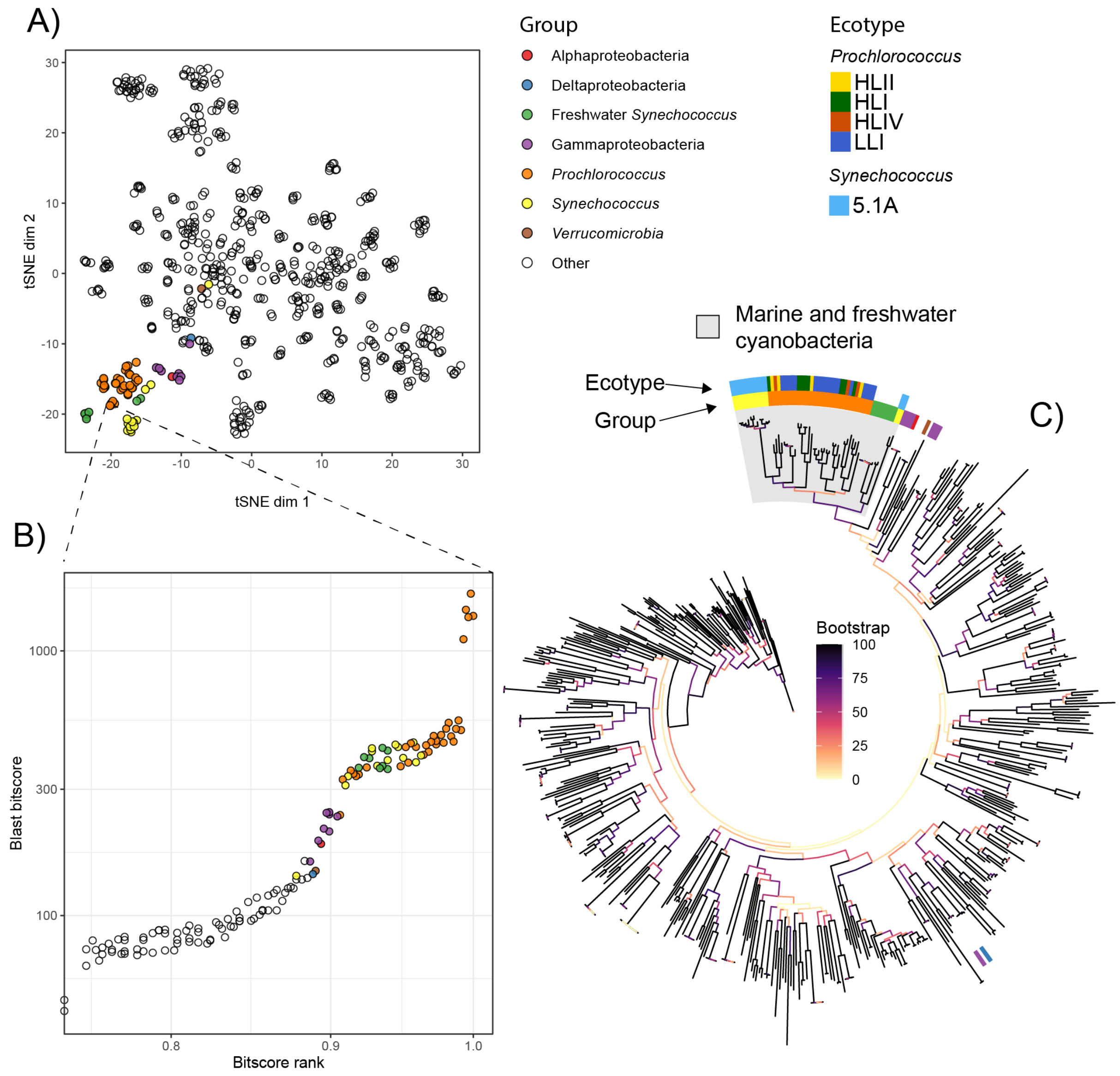

**Supplementary Figure S2.** Comparing the sequence similarity of the picocyanobacterial TonB-dependent transporters (TBDT) with TBDTs from other functional/taxonomic groups in MARMICRODB
**A)** t-Distributed Stochastic Neighbor Embedding (tSNE) dimensionality reduction of pairwise sequence similarity score between 625 tonB dependent transporter amino acid sequences with a pairwise blastp e value < 1e-10 to any marine picocyanobacterial TBDT. Sequences that share greater amino acid similarity cluster more closely together. The colored subset represents all sequences with an e value < 1e-40 to a marine picocyanobacterial TBDT, and they are colored by their taxonomic origin. **B)** Focused view of the sequence similarity scores of the TBDT from MIT0917 and the 624 other TBDTs. Each point is a sequence comparison to the MIT0917 sequence, and the vertical axis shows the blastp bitscore of the comparison. The comparisons are bitscore rank-ordered (x-axis), showing only those from the top 25%. **C)** Unrooted maximum likelihood phylogenetic tree (RAxML) of the 625 TBDTs from A) and B). Branches are colored by bootstrap score out of 300 bootstraps. The inner ring highlights the same highlighted sequences from A) and B), and the outer ring categorizes sequences by picocyanobacterial clade.

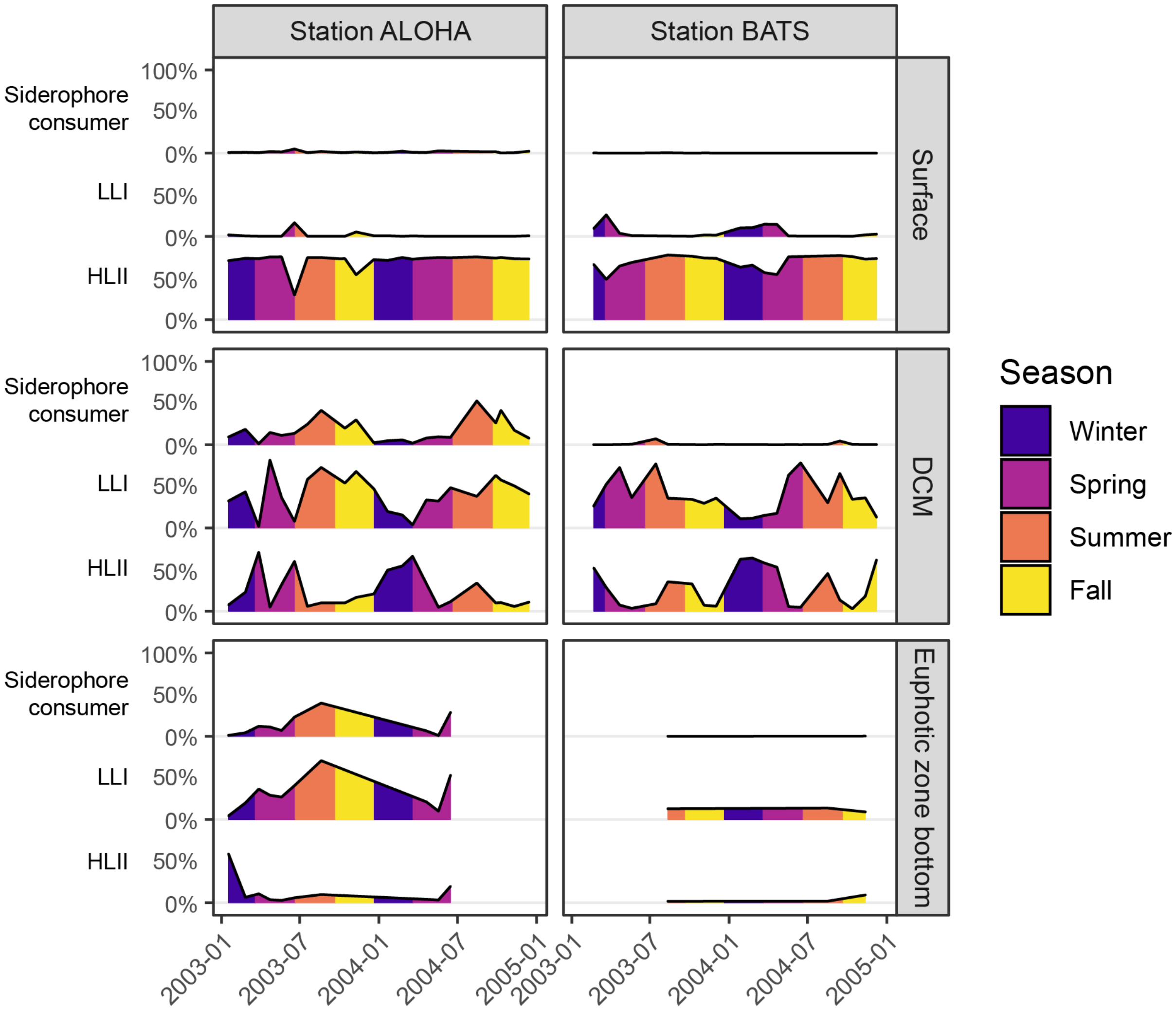

**Supplementary Figure S3.** Seasonal dynamics of siderophore consumers in the N. Pacific (Station ALOHA) and N. Atlantic (Station BATS)

Relative abundance (%) of *Prochlorococcus* siderophore consumers (top), LLI clade (middle), and HLII clade (bottom) at the surface, deep chlorophyll maximum layer, and the bottom of the euphotic zone at HOT and BATS from 2003 to 2005. Area plots are colored by the season when samples were collected.

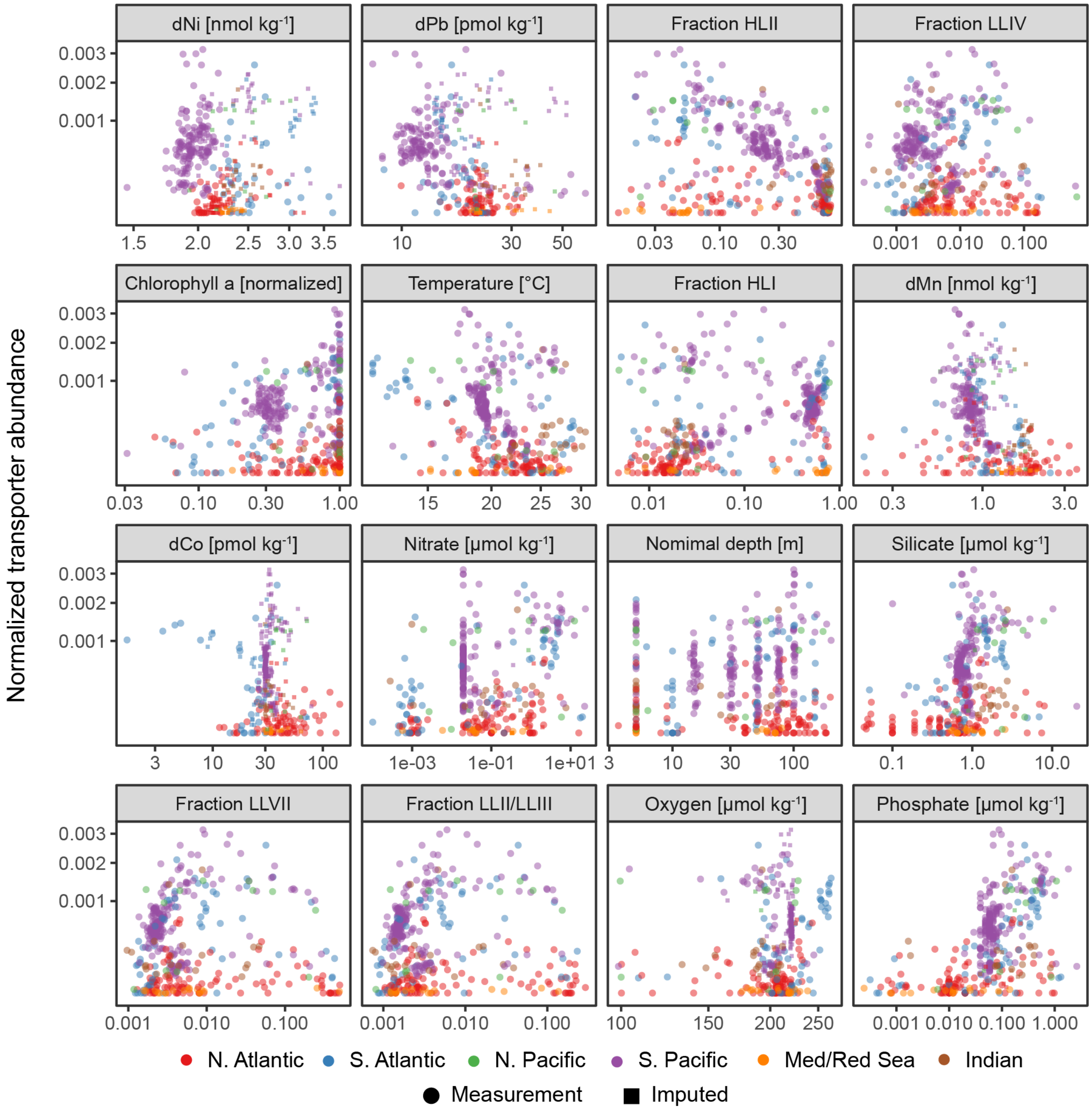

**Supplementary Figure S4.** Other predictive variables of *Prochlorococcus* siderophore consumers
Normalized abundance of *Prochlorococcus* siderophore transporters plotted against the remaining environmental variables with the greatest predictive contribution to the random forest model. Each point is a metagenomic observation colored by ocean basin. Small squares represent samples with missing data that was imputed for each variable (see methods). Dissolved Fe climatological means are from the MIT Darwin model. The remaining measurements are *in situ* chemical, biological, or hydrographic measurements from GEOTRACES and Tara Oceans.

### Supplementary Tables:

**Supplementary Table S1.** Regression of dFe, Fe-binding ligands, and siderophores

**A) Model summary** (GLM Gaussian)

| Measurement | Variable | Estimate | Std. Error | t value | p-value |
|---|---|---|---|---|---|
| dFe [nmol kg$^{-1}$] | (Intercept) | -1.0862 | 0.1959 | -5.55 | 8.60e-07 |
| dFe [nmol kg$^{-1}$] | $Ocean_{N. Pacific}$ | -1.0749 | 0.2287 | -4.70 | 1.78e-05 |
| dFe [nmol kg$^{-1}$] | $Env_{Surface}$ | 0.2771 | 0.3158 | 0.88 | 0.3840 |
| dFe [nmol kg$^{-1}$] | $ocean_{N. Pacific}$:$Env_{Surface}$ | 0.6793 | 0.3556 | 1.91 | 0.0613 |
| | AIC=103.592; BIC=113.980; $R^2$ = 0.55; RMSE=0.535; Dispersion = 0.306 | | | | |
| $L_1$ [nmol kg$^{-1}$] | (Intercept) | 0.3031 | 0.1136 | 2.67 | 0.0110 |
| $L_1$ [nmol kg$^{-1}$] | $Ocean_{N. Pacific}$ | -1.0062 | 0.1444 | -6.97 | 2.36e-08 |
| $L_1$ [nmol kg$^{-1}$] | $Env_{Surface}$ | -0.1243 | 0.1832 | -0.68 | 0.5015 |
| $L_1$ [nmol kg$^{-1}$] | $ocean_{N. Pacific}$:$Env_{Surface}$ | 0.8143 | 0.2181 | 3.73 | 0.0006 |
| | AIC=30.192; BIC=38.998; $R^2$ = 0.65; RMSE=0.306; Dispersion = 0.103 | | | | |

**B) Model contrasts**

| Measurement | Ref. level | Contrast | Estimate | Std. Error | z-ratio | p-value |
|---|---|---|---|---|---|---|
| dFe [nmol kg$^{-1}$] | DCM | N. Pacific - N. Atlantic | -1.0749 | 0.2287 | -4.700 | <.0001 |
| dFe [nmol kg$^{-1}$] | Surface | N. Pacific - N. Atlantic | -0.3956 | 0.2723 | -1.453 | 0.5850 |
| dFe [nmol kg$^{-1}$] | N. Atlantic | Surface - DCM | 0.2771 | 0.3158 | 0.878 | 1.0000 |
| dFe [nmol kg$^{-1}$] | N. Pacific | Surface - DCM | 0.9564 | 0.1635 | 5.849 | <.0001 |
| $L_1$ [nmol kg$^{-1}$] | DCM | N. Pacific - N. Atlantic | -1.0062 | 0.1444 | -6.969 | <.0001 |
| $L_1$ [nmol kg$^{-1}$] | Surface | N. Pacific - N. Atlantic | -0.1919 | 0.1635 | -1.174 | 0.9615 |
| $L_1$ [nmol kg$^{-1}$] | N. Atlantic | Surface - DCM | -0.1243 | 0.1832 | -0.678 | 1.0000 |
| $L_1$ [nmol kg$^{-1}$] | N. Pacific | Surface - DCM | 0.6900 | 0.1184 | 5.829 | <.0001 |

**A)** Regression summary for dFe and strong Fe-binding ligands ($L_1$) as a function of Ocean and depth (surface or DCM). Response error distributions was Gaussian with identity-link. **B)** Pairwise model contrasts. P value adjustment: Bonferroni method for 4 tests. Results are given on the log (not the response) scale.

**Supplementary Table S2.** Beta regression of siderophore transporter abundance

**A) Model summary** (mean model with logit link)

| Coefficient | Estimate | Std. Error | z value | p-value |
|---|---|---|---|---|
| (Intercept) | -8.70782 | 0.16019 | -54.359 | < 2e-16 |
| LLI abund.$_{mid}$ | 0.18176 | 0.11633 | 1.562 | 0.11817 |
| LLI abund.$_{high}$ | 0.25708 | 0.11523 | 2.231 | 0.02569 |
| Nitrite conc.$_{mid}$ | 0.68633 | 0.30074 | 2.282 | 0.02248 |
| Nitrite conc.$_{high}$ | 0.88253 | 0.17135 | 5.150 | 2.6e-07 |
| DCM Depth$_{mid}$ | -0.22344 | 0.19631 | -1.138 | 0.25504 |
| DCM Depth$_{high}$ | 0.06889 | 0.20265 | 0.340 | 0.73390 |
| Nitrite$_{mid}$:DCM$_{mid}$ | 0.73436 | 0.35598 | 2.063 | 0.03912 |
| Nitrite$_{high}$:DCM$_{mid}$ | 0.66712 | 0.25025 | 2.666 | 0.00768 |
| Nitrite$_{mid}$:DCM$_{high}$ | 0.53029 | 0.35621 | 1.489 | 0.13657 |
| Nitrite$_{high}$:DCM$_{high}$ | 0.60225 | 0.27292 | 2.207 | 0.02733 |

Pseudo $R^2$ = 0.40; Log Likelihood = 2948.85; Df = 12; Num. obs. = 407

**B) Joint tests of the terms in the model**

| Model Term | Df1 | Df2 | F ratio | p-value |
|---|---|---|---|---|
| LLI abund. | 2 | 12 | 2.87 | 0.0528 |
| Nitrite conc. | 2 | 12 | 56.28 | <.0001 |
| DCM Depth | 2 | 12 | 8.31 | 0.0002 |
| Nitrite conc:DCM Depth | 4 | 12 | 4.45 | 0.0014 |

**C) Model contrasts**

| DCM Depth | LLI abund. | Contrast Nitrite conc. | Estimate | Std. Error | z ratio | p-value |
|---|---|---|---|---|---|---|
| mid | low | mid - low | 4.15e-04 | 7.71e-05 | 5.38 | 7.85e-07 |
| mid | mid | mid - low | 4.97e-04 | 7.94e-05 | 6.27 | 3.24e-09 |
| mid | high | mid - low | 5.36e-04 | 9.37e-05 | 5.73 | 9.32e-08 |
| high | low | mid - low | 4.20e-04 | 7.69e-05 | 5.47 | 6.99e-07 |
| high | mid | mid - low | 5.04e-04 | 7.33e-05 | 6.88 | 4.46e-11 |
| high | high | mid - low | 5.43e-04 | 9.41e-05 | 5.77 | 7.11e-08 |

**A)** Beta regression summary for siderophore transporter relative abundance as function of LLI abundance [%], $NO_2$ concentration [μmol $kg^{-1}$], and DCM depth [m]. Response error distributions was Beta with logit-link. Estimated coefficients are on the logit scale. **B)** Interaction contrasts for all effects in the model compiled into a Type-III-ANOVA-like table. **C)** Pairwise model contrasts. For brevity only contrasts with $P \leq 0.05$ are included. P value adjustment: mvt method for 54 tests. Tests are performed on the logit scale.
